# Overexpression of the vascular brassinosteroid receptor BRL3 confers drought resistance without penalizing plant growth

**DOI:** 10.1101/318287

**Authors:** Norma Fàbregas, Fidel Lozano-Elena, David Blasco-Escámez, Takayuki Tohge, Cristina Martínez-Andújar, Alfonso Albacete, Sonia Osorio, Mariana Bustamante, José Luis Riechmann, Takahito Nomura, Takao Yokota, Ana Conesa, Francisco Pérez Alfocea, Alisdair R. Fernie, Ana I. Caño-Delgado

## Abstract

Drought represents a major threat to food security. Mechanistic data describing plant responses to drought have been studied extensively and genes conferring drought resistance have been introduced into crop plants. However, plants with enhanced drought resistance usually display lower growth, highlighting the need for strategies to uncouple drought resistance from growth. Here, we show that overexpression of BRL3, a vascular-enriched member of the brassinosteroid receptor family, can confer drought stress tolerance in *Arabidopsis*. Whereas loss-of-function mutations in the ubiquitously expressed BRI1 receptor leads to drought resistance at the expense of growth, overexpression of BRL3 receptor confers drought tolerance without penalizing overall growth. Systematic analyses reveal that upon drought stress, increased BRL3 triggers the accumulation of osmoprotectant metabolites including proline and sugars. Transcriptomic analysis suggests that this results from differential expression of genes in the vascular tissues. Altogether, this data suggests that manipulating BRL3 expression could be used to engineer drought tolerant crops.

## INTRODUCTION

Drought is responsible for at least 40% of crop losses worldwide and this proportion is dramatically increasing due to climate change ^1^. Understanding cellular responses to drought stress represents the first step toward the development of better-adapted crops, something which is a great challenge for the field of plant biotechnology ^2^. Classical approaches aimed at examining how plants cope with limited water led to the identification of regulators involved in the signal transduction cascades of the abscisic acid (ABA)-dependent and ABA-independent pathways ^3^. Adaptation to drought stress has been associated with the presence of proteins that protect cells from dehydration, such as late-embryogenesis–abundant (LEA) proteins, osmoprotectants and detoxification enzymes ^4,5^. These studies provided deep insights into the molecular mechanisms underlying abiotic stress ^2^, showing that drought resistance is a complex trait simultaneously controlled by many genes. While genetic approaches have succeeded in conferring stress resistance to plants, this generally comes at the cost of reduced growth ^6,7^. Therefore, understanding how cellular growth is coupled to drought stress responses is essential for engineering plants with improved growth in rain-fed environments.

Receptor-like kinases (RLKs) play an important role in optimizing plant responses to stress ^8,9^. Brassinosteroid (BR) hormones directly bind to BRI1 (BR-INSENSITIVE 1) leucine-rich repeat (LRR)-RLK family members on the plasma membrane ^10-14^. Ligand perception triggers BRI1 to interact with the co-receptor BAK1 (BRI1 ASSOCIATED RECEPTOR KINASE 1) ^15-17^, which is essential for early BR signaling events ^18^. This BRI1-BAK1 heterodimerization initiates a signaling cascade of phosphorylation events that control the expression of multiple BR-regulated genes mainly via the BRI1-EMS-SUPPRESSOR1 (BES1) and BRASSINAZOLE RESISTANT1 (BZR1) transcription factors ^19-21^.

Although BRs modulate multiple developmental and environmental stress responses in plants, the exact role of BRs under stress conditions remains controversial. Whereas the exogenous application of BRs and the overexpression of the BR biosynthetic enzyme DWF4 both confer increased plant adaptation to drought stress ^22-24^, suppression of the BRI1 receptor also results in drought-resistant phenotypes ^25,26^. Intriguingly, ABA signaling inhibits the BR signaling pathway after BR perception, and crosstalk between the two pathways upstream of the BIN2 (BRASSINOSTEROID-INSENSITIVE 2) kinase has been reported ^27,28^. Further crosstalk has been described downstream mediated by the overlapping transcriptional control of multiple BR- and ABA-regulated genes ^29,30^ such as *RESPONSE TO DESICCATION 26* (*RD26*) ^26^.

Recently, greater attention is being placed on the spatial regulation of hormonal signaling pathways in attempt to further understand the coordination of plant growth and stress responses ^26,31-34^. For instance, while the BRI1 receptor is widely localized in many tissues ^35^, the BRI1-LIKE receptor homologues BRL1 and BRL3 signal from the innermost tissues of the plant and thereby contribute to vascular development ^12,33,36^. BR receptor complexes are formed by different combinations of BRI1-like LRR-RLKs with the BAK1 co-receptor in the plasma membrane ^33^. Despite BRI1 being a central player in plant growth and adaptation to abiotic stress ^26,37,38^, the functional relevance of vascular BRL1 and BRL3 is only just beginning to be explored ^33,39^. For example, in previous proteomic approaches we found abiotic stress-related proteins within BRL3 signalosome complexes ^33^, but the exact role of the BRL3 pathway in drought remains elusive.

Here, we show that knocking out or overexpressing different BR receptors modulate multiple drought stress-related traits in both the roots and shoots. While the traits controlled by the BRI1 pathway are intimately linked to growth arrest, we found that overexpressing the vascular-enriched BRL3 receptors can confer drought resistance without penalizing overall plant growth. Moreover, metabolite profiling revealed that the overexpression of the BRL3 receptor triggers the production of an osmoprotectant signature (i.e., proline, trehalose, sucrose, and raffinose family oligosaccharides) in the plant and the specific accumulation of the osmoprotectant metabolites in the roots during periods of drought. Subsequent transcriptomic profiling showed that this metabolite signature is transcriptionally regulated by the BRL3 pathway in response to drought. An enrichment of deregulated genes in root vascular tissues, especially in the phloem, further supports a preferential accumulation of osmoprotectant metabolites to the root. Overall, this study demonstrates that overexpression of the BRL3 receptor boosts the accumulation of sugar and osmoprotectant metabolites in the root and overcomes drought-associated growth arrest, thereby uncovering a strategy to protect crops against drought.

## RESULTS

### BR receptors control osmotic stress sensitivity in the root

To determine the contribution of the BR complexes in the response to drought, we performed a comprehensive characterization of different combinations of mutants of all the BR receptors and the BAK1 co-receptor. For each combination, we first analyzed primary root growth (Fig. 1a). As previously described ^17,33,40^, seven-day-old roots of *bak1*, *brl1brl3bak1*, *bri1*, and *bri1brl1brl3* displayed shorter roots than the Col-0 wild type (WT). We also found that the primary roots of the quadruple mutant *bri1brl1brl3bak1* (hereafter *quad*) were the shortest and the most insensitive to BRs (Fig. 1a,b and Supplementary Fig. 1). Conversely, plants overexpressing BRL3 (*35S:BRL3-GFP*, hereafter *BRL3ox*) not only exhibited longer roots than WT (Fig. 1a,b) but also showed increased receptor levels in root vascular tissues ^33^ (Supplementary Fig. 2). These results agree with the previously reported role of BR receptors in promoting root growth ^40,41^. We then subjected Arabidopsis seedlings to osmotic stress by transferring them to sorbitol-containing media and subsequently quantified the level of inhibition of root growth in sorbitol relative to control conditions (see Methods). A significantly lower level of relative root growth inhibition mediated by osmotic stress was observed in *bri1* (27%), *bri1brl1brl3* (28%) and *quad* (27%) mutants compared to the WT (39%; Fig. 1a,b). In contrast, no differences were found in *brlbrl3* and *brl1brl3bak1* root growth inhibition when compared to WT (Fig. 1a,b). Similarly, the roots of *BRL3ox* plants were like those of WT in terms of relative root growth inhibition (Fig. 1a,b).

**Figure 1.**
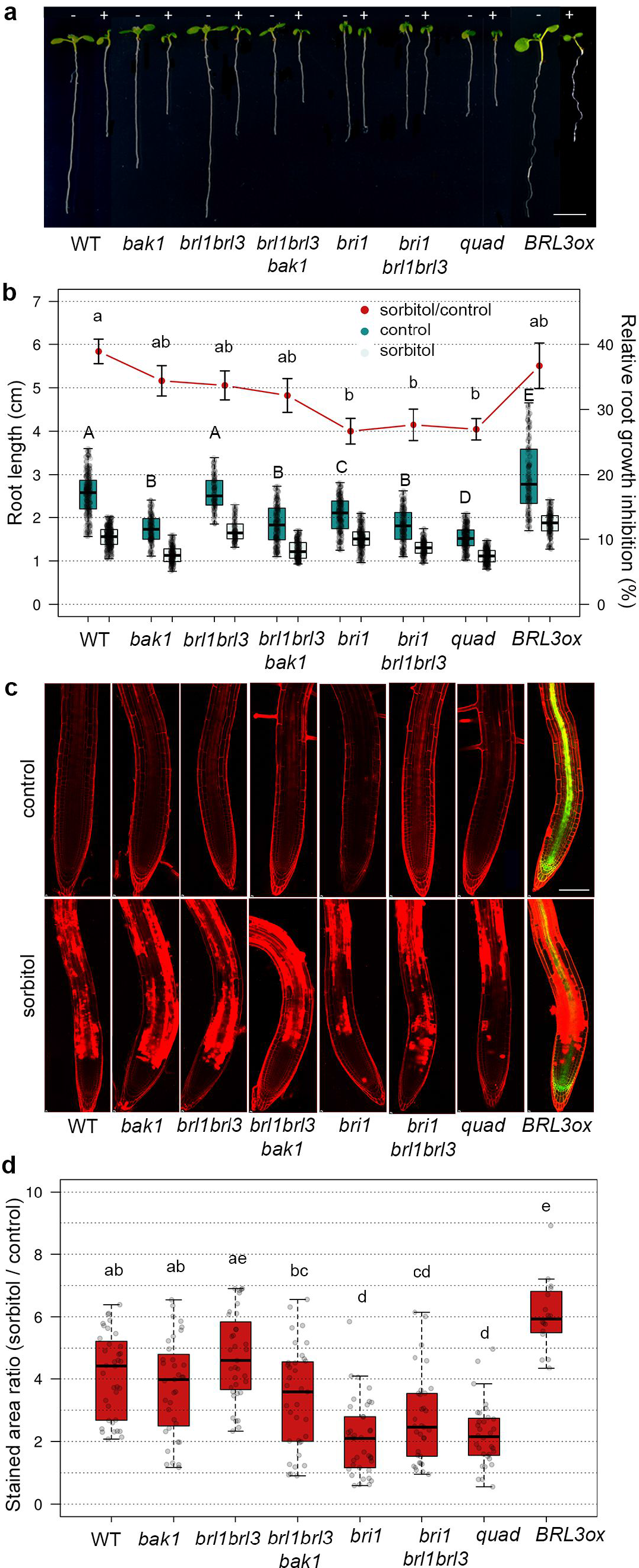
BR perception mutant roots are less sensitive to osmotic stress. (**a**) Seven-day-old roots of WT, BR mutants *bak1*, *brl1brl3, brl1brl3bak1*, *bri1, bri1brl1brl3*, and *bri1brl1brl3bak1* (*quad*), and BR overexpressor line *35S:BRL3-GFP (BRL3ox)* grown in control (-) or 270 mM sorbitol (+) conditions. Scale bar: 0.5 cm. (**b**) Boxplots depict the distribution of seven-day-old root lengths in control (dark green) or sorbitol (light green) conditions. Red line depicts relative root growth inhibition upon stress (ratio sorbitol/control +/-s.e.m.). Data from five independent biological replicates (n>150). Different letters represent significant differences (p-value<0.05) in an ANOVA plus Tukey’s HSD test. (**c**) Four-day-old roots stained with propidium iodide (PI, red) after 24 h in control (top) or sorbitol (bottom) media. Green channel (GFP) shows the BRL3 membrane protein receptor under the 35SCaMV constitutive promoter localizing to the vascular tissues in primary roots. Scale bar: 100 μmm. (**d**) Quantification of cell death in sorbitol-treated root tips. Boxplots represent the relative PI staining (sorbitol/control) for each genotype. Averages from five independent biological replicates (n>31). Different letters represent significant differences (p-value<0.05) in an ANOVA plus Tukey’s HSD test. Boxplots represent the median and interquartile range (IQR). Whiskers depict Q1-1.5*IQR and Q3+1.5*IQR and points experimental observation.

Previous experimental evidences unveiled that water stress-induced cell death in Arabidopsis roots is localized and occurs via programmed cell death (PCD) ^42^. As shown by the incorporation of propidium iodine (PI) into the nuclei (Fig. 1c,d), a short period of osmotic stress (24h) caused cell death in the elongation zone of WT roots. In comparison, a reduced amount of cell death was observed in the roots of *bri1, bri1brl1brl3* and *quad* mutants (Fig. 1c,d), thereby indicating less sensitivity towards osmotic stress. Conversely, plants with increased levels of BRL3 showed a massive amount of cell death in root tips compared to WT, indicating an increased sensitivity to short osmotic stress (Fig 1c,d). These results point towards a role for BR receptors in triggering osmotic stress responses in the plant root.

Since root hydrotropism represents a key feature for adaptation to environments scarce in water ^43^, we investigated the capacity of roots to escape imposed osmotic stress by bending towards water-available media (Fig. 2a). We found that BR receptor loss-of-function mutants had reduced hydrotropic responses compared to WT plants. For instance, while no significant differences were found under control conditions (mock) (Supplementary Fig. 3), the roots of BR receptor mutants grew straighter than WT roots towards sorbitol-containing media (Fig. 2a-c). Interestingly, *brl1brl3bak1* mutants were the least sensitive to osmotic stress in terms of hydrotropism, showing lower root curvature angles than the *quad* roots (Fig. 2b). Consistently, compared to WT roots, an enhanced hydrotropic response was observed in *BRL3ox* (Fig. 2a-c). Furthermore, exogenous application of the BR synthesis inhibitor brassinazole ^44^ reverted the hydrotropic response of WT roots (Supplementary Fig. 3). For better visualization, we generated a drought multi-trait matrix for all the BR receptor mutants analyzed in this study (Fig. 2d; Supplementary Table 1). From this matrix, it can be seen that overexpression or mutation of BRL3/BRL1/BAK1 receptors in the vascular tissues alters drought-response related traits.

**Figure 2.**
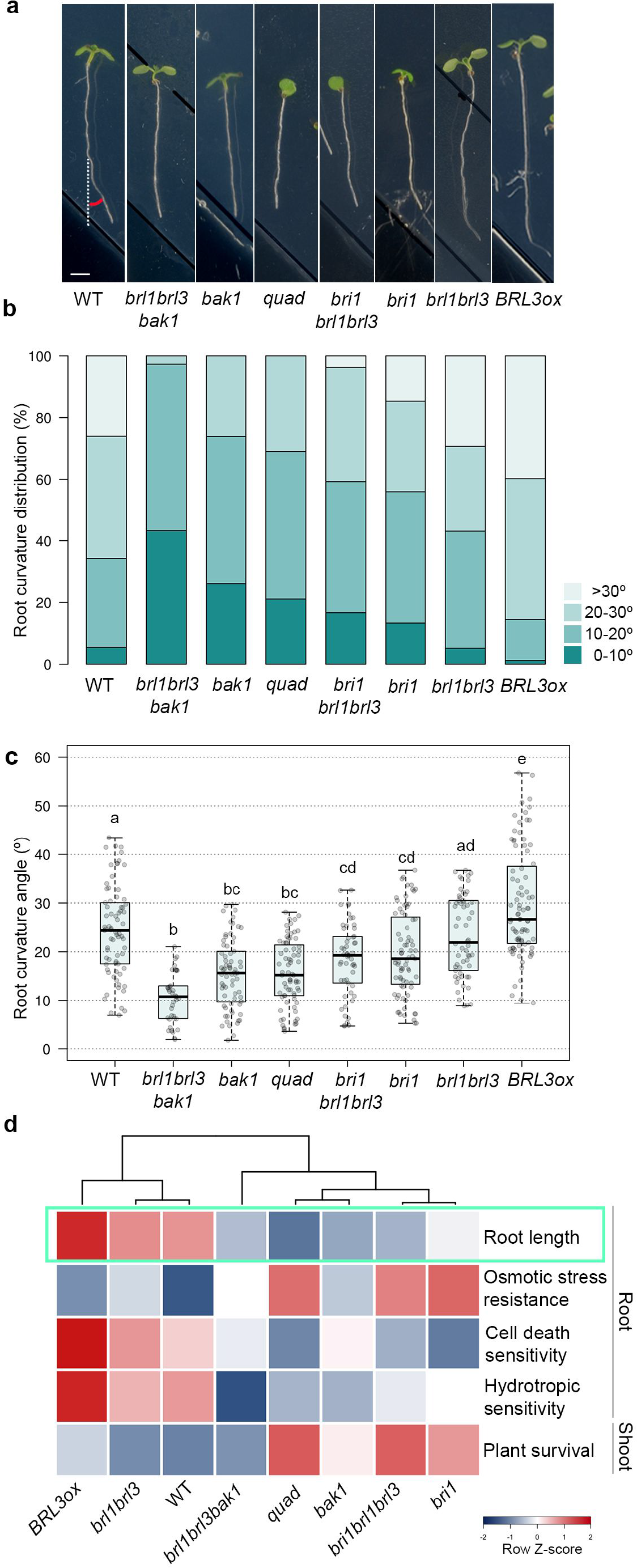
Overexpression of the BRL3 receptor promotes root hydrotropism. (**a**) Root curvature (hydrotropic response) in seven-day-old roots after 24 h of sorbitol-induced osmotic stress (270 mM). Scale bar: 0.2 cm. (**b**) Discrete distribution of root hydrotropic curvature angles in the different genotypes. Lightest green depicts roots curved between 0° and 10°, light green between 10° and 20°, dark grey between 20° and 30°, and darkest green depicts roots that have a curvature of more than 30° as indicated in the color legend. (**c**) Continuous distribution of root curvature angles. Different letters indicate a significant difference (p-value<0.05) in a one-way ANOVA test plus Tukey’s HSD test. Boxplot represent the median and interquartile range (IQR). Whiskers depict Q1-1.5*IQR and Q3+1.5*IQR and points experimental observations. Data from four independent biological replicates (n>50). (**d**) Stress traits matrix for all physiological assays performed on the roots and shoots of WT, BR loss-of-function mutants and *BRL3ox*. Root growth in control conditions is highlighted in green. Color bar depicts values for scaled data.

### *BRL3ox* confers drought resistance without penalizing growth

To investigate if the impaired responses to abiotic stress observed in root seedling were preserved in mature plants, we next analyzed the phenotypes of plants exposed to severe drought. After 12 days of withholding water, dramatic symptoms of drought stress were observed in WT, *brl1brl3* and *brl1brl3bak1* mutants. In contrast, other BR mutants showed a remarkable degree of drought resistance. In particular, *bak1*, *bri1*, *bri1brl1brl3*, and *quad* mutant plants were the most resistant to the severe water-withholding regime (Fig. 3a). As these mutants exhibited some degree of dwarfism (Fig. 3a), we confirmed their resistance to drought by examining their survival rates after re-watering (Fig. 3b). To correct for the delayed growth seen in BR-deficient mutants, plants were submitted to a time course of drought stress in which water use, photosynthesis and transpiration parameters were monitored under similar relative soil water content (Fig. 3c-e). The WT plants took just 9 days to use 70% of the available water (field capacity) during the drought period (Fig. 3c). In comparison, BR loss-of-function mutant plants *bri1*, *bri1brl1brl3* and *quad* took 15 days. All subsequent measurements were done at the same soil water content for each genotype. We found that the relative water content (RWC) in WT plants was reduced during drought, while RWC in BR mutant leaves remained as in well-watered conditions (Fig. 3d). In addition, compared to WT plants, BR mutants sustained higher levels of photosynthesis and transpiration during the drought period (Fig. 3e and Supplementary Fig. 4). Altogether our results indicate that the dwarf BR receptor mutant plants are more resistant while consuming less water, likely through avoiding the effects of drought (Fig. 3f).

**Figure 3.**
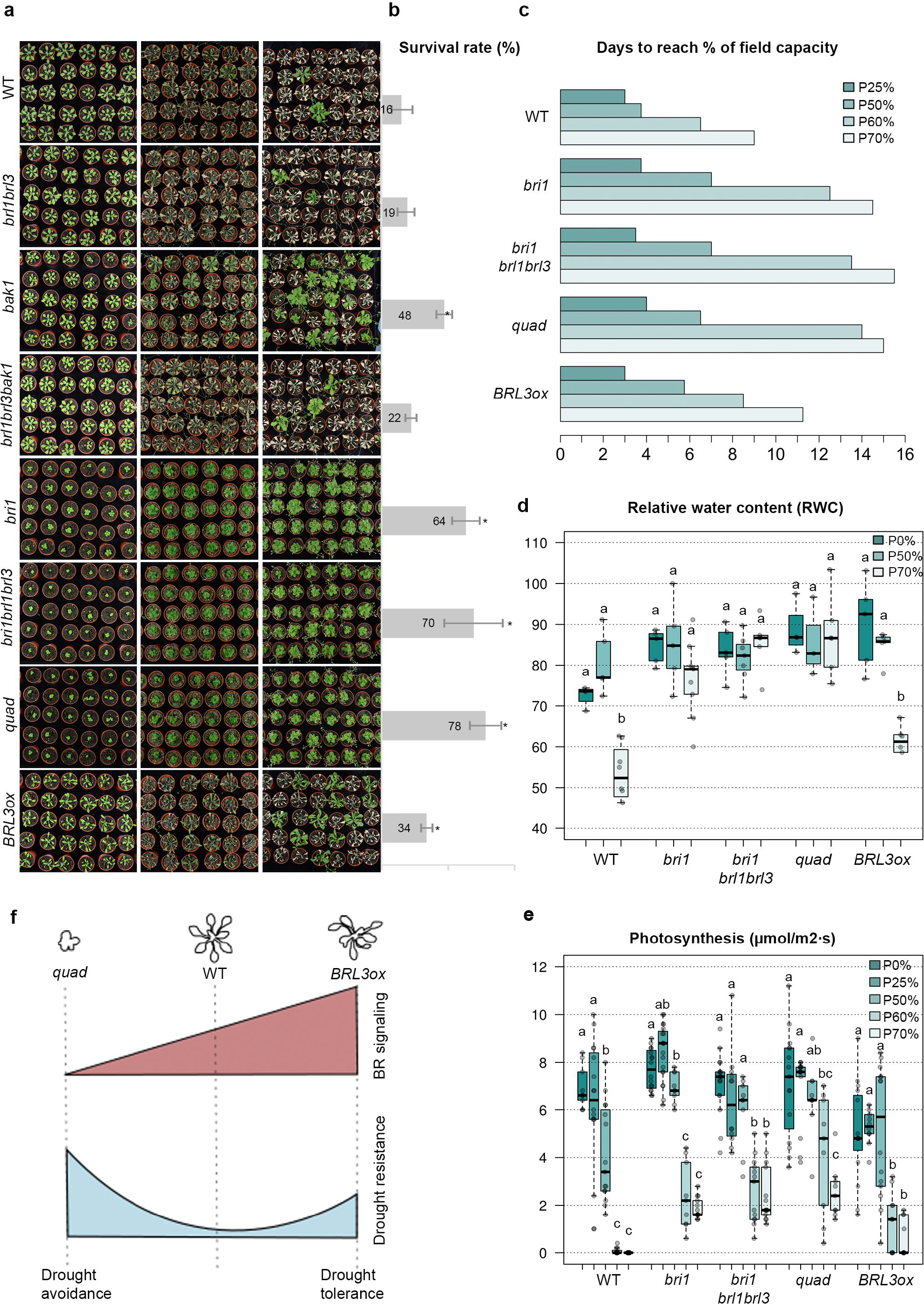
BRL3 overexpression confers drought tolerance. (**a**) From top to bottom, three-week-old plant rosette phenotypes of WT, *brl1brl3, bak1*, *brl1brl3bak1*, *bri1, bri1brl1brl3, quad* and *BRL3ox* grown in well-watered conditions (left column), after 12 days of drought stress (middle column) and after 7 days of re-watering (right column). (**b**) Plant survival rates after 7 days of re-watering. Averages of five independent biological replicates +/- s.e.m. (n>140). Asterisk indicates a significant difference (p-value<0.05) in a chi-squared test for survival ratios compared to WT. (**c**) Bar plot shows the days needed to reach different percentages of the soil field capacity for each genotype used in the study. **(d)** Relative water content (RWC) of mature rosettes at 0% (field capacity), 50% and 70% soil water loss. **(e**) Photosynthesis efficiency (µmol/m2*s) at different percentages of soil water loss. (**d, e**) Boxplot represent the median and interquartile range (IQR). Whiskers depict Q1-1.5*IQR and Q3+1.5*IQR and points experimental observations (n=6). Different letters depict significant differences within each genotype in a one-way ANOVA plus a Tukey’s HSD test. (**f**) Schematic representation of BR signaling levels, adult plant size and drought resistance. Loss-of-function mutants passively avoid stress (drought avoidance), whereas plants with increased levels of BRL3 act actively to avoid drought stress (drought tolerance).

Strikingly, we found that *BRL3ox* plants were more resistant than WT plants to severe drought stress as shown by increased survival rates (Fig. 3a,b). Plants with increased BRL3 receptors showed reduction of RWC during drought similarly to WT plants (Fig. 3d). Interestingly the rate of photosynthesis was lower in *BRL3ox* compared to WT at basal conditions, but together with transpiration, was more stable than in WT plants during the drought period (Fig. 3,e and Supplementary Fig. 4). This indicates that *BRL3ox* plants are healthier than WT under the same water consumption conditions. These results suggest that the BRL3 overexpression actively promotes drought tolerance without penalizing plant growth (Fig. 3f).

### *BRL3ox* plants accumulate osmoprotectant metabolites

To further investigate the cause behind drought tolerance conferred by BRL3 overexpression, we performed metabolite profiling of *BRL3ox* plants and compared it to the profile of WT and *quad* plants in a time course drought experiment. Roots were separated from shoots to address possible changes in metabolite accumulation from source to sink tissues. The complete metabolic fingerprints are provided in Supplementary Fig. 5 and 6 and Supplementary Data 1 and 2. Metabolite profiling of mature *BRL3ox* plants grown in control conditions (time 0) revealed an increment in the production of osmoprotectant metabolites. Both shoots (Fig. 4a) and roots (Fig. 4b) of the *BRL3ox* plants exhibited metabolic signatures enriched in proline and sugars, metabolites which have previously been reported to confer resistance to drought ^45-47^. This suggests that the BRL3 receptor promotes priming ^48^. Importantly, the levels of these metabolites were lower in *quad* mutant plants (Fig. 4a,c).

**Figure 4.**
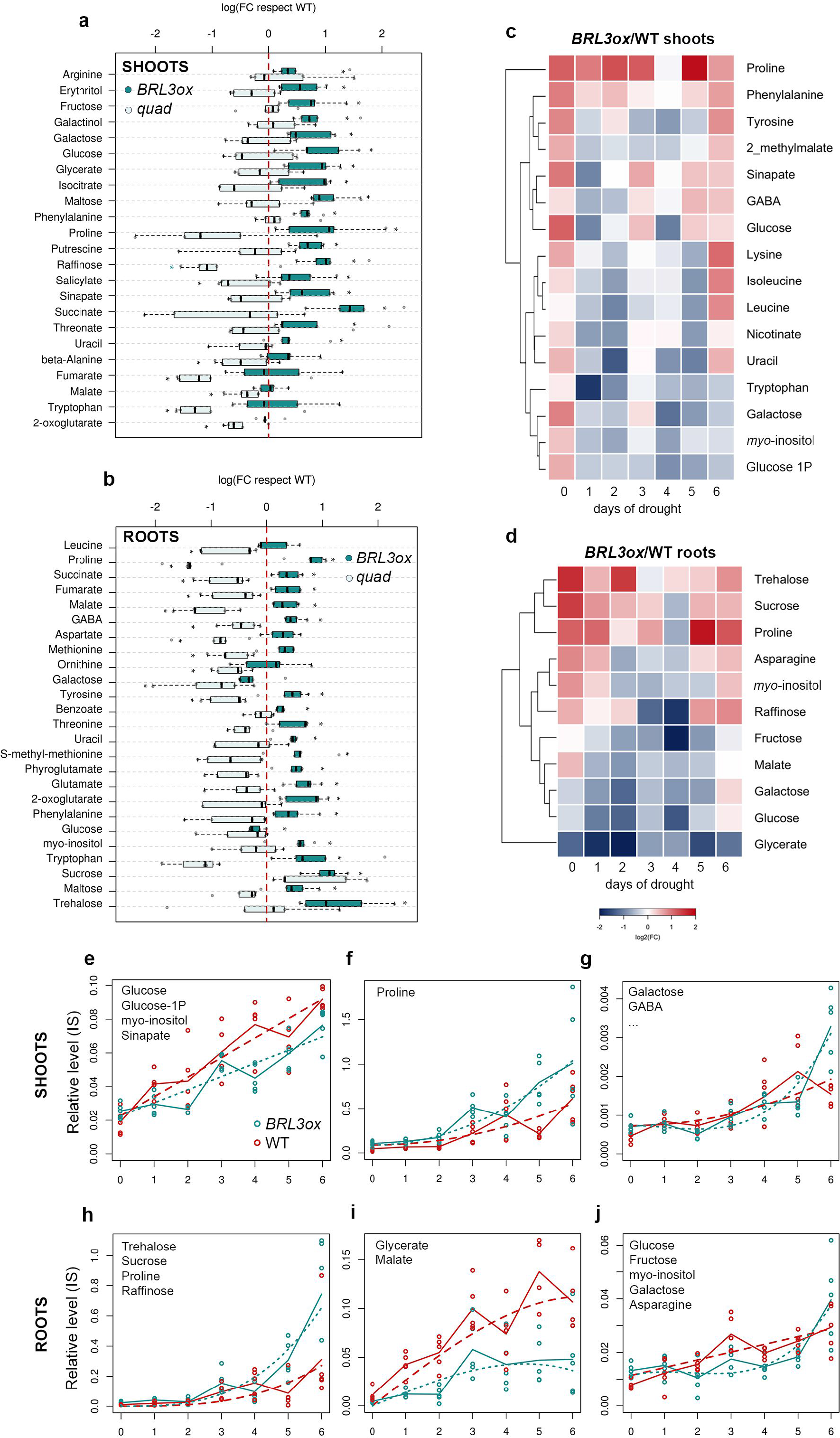
BRL3 overexpression plants show a primed metabolic signature. (**a**) Metabolites differentially accumulated in *BRL3ox* (dark green) *or quad* (light green) shoots relative to WT at basal conditions. (**b**) Metabolites differentially accumulated in *BRL3ox* (dark green) *or quad* (light green) roots relative to WT at basal conditions. (**a, b**) Boxplot represent the median and interquartile range (IQR). Whiskers depict Q1-1.5*IQR and Q3+1.5*IQR and points experimental observations (n=5). Asterisks denote statistical differences in a two-tailed t-test (p-value < 0.05) for raw data comparisons *BRL3ox* vs. WT (panel right side) or *quad* (panel left side). (**c**) Metabolites following differential dynamics between *BRL3ox* and WT shoots along the drought time course. (**d**) Metabolites following differential dynamics between *BRL3ox* and WT roots along the drought time course. (**c, d**) Heatmap represents the log2 ratio of *BRL3ox*/WT. (**e-j**) Clustering of the dynamics of relative metabolite levels along the drought time course in shoots and roots. Solid lines show the actual metabolic profile (averages) of the representative metabolite for each cluster while dashed lines represent the polynomial curve that best fit the profile. Statistical significance was evaluated with the maSigPro package. (**e**) Metabolites following a linear increase during drought in shoots include Glucose, Glucose-1P, *myo*-inositol, and Sinapate. (**f**) Proline follows a steeper exponential increase in *BRL3ox* shoots. (**g**) Metabolites following an exponential increase in *BRL3ox* shoots but nearly a linear increase in WT include galactose, GABA, phenylalanine, tyrosine, 2-methylmalate, lysine, isoleucine, leucine, nicotinate, uracil, and tryptophan. (**h**) Metabolites following a steeper exponential increase in *BRL3ox* roots include trehalose, sucrose, proline and raffinose. (**i**) Metabolites following a reduced linear increase until a certain maximum in *BRL3ox* roots include glycerate and malate. (**j**) Metabolites following an exponential increase in *BRL3ox* roots but a linear increase in WT include glucose, fructose, *myo-*inositol, galactose, and asparagine.

Compared to WT, sugars including fructose, glucose, galactinol, galactose, maltose, and raffinose overaccumulated in the shoots of *BRL3ox* (Fig. 4a). Conversely, whereas glucose levels were lower in the roots, sucrose, trehalose, *myo*-inositol, and maltose appeared to accumulate here (Fig. 4b) suggesting that the BRL3 pathway promotes sugar accumulation preferentially in the roots. We then analyzed the dynamics of each metabolite in response to drought (see Methods). In this time course, a rapid accumulation of osmoprotectant metabolites was observed in *BRL3ox* plants (Fig. 4c,d). In the shoots, proline showed the highest levels respect WT along the entire drought time course (Fig. 4c,f). In contrast, glucose, galactose and *myo*-inositol increased at similar or slightly lower rates than in the shoots of WT plant (Fig. 4c,e,g). However, in roots, an accumulation of trehalose, sucrose, proline, and raffinose was observed in *BRL3ox* mutants subjected to drought stress (Fig. 4d), and this accumulation showed steeper exponential dynamics than in WT plants (Fig. 4h). Additionally, glucose, galactose, fructose, and *myo*-inositol linearly increased in WT roots but exponentially increased in *BRL3ox* roots (Fig. 4j). Interestingly, throughout this time course, the levels of these metabolites were lower in the *quad* mutant plants compared to in WT (Supplementary Fig. 7). Altogether, these findings uncover a key role for BR receptors in promoting sugar metabolism, and support the idea that BRL3 triggers the accumulation of osmoprotectant metabolites in the root to promote growth during periods of drought.

### Transcriptional control of metabolite production in *BRL3ox*

We next investigated whether metabolic pathways are transcriptionally regulated in *BRL3ox* roots. RNAseq of *BRL3ox* roots revealed 759 deregulated genes at basal conditions (214 upregulated and 545 downregulated; FC>1.5, FDR<0.05; Supplementary Data 3) and 1,068 deregulated genes in drought conditions (378 upregulated and 690 downregulated; FC>1.5, FDR<0.05; Supplementary Data 4). In control conditions, a high proportion of the deregulated genes belonged to the response to water stress, oxygen-containing compounds (ROS) and response to ABA GO categories (Fig. 5a, 5c, Supplementary Data 5 and 6). We next deployed the genes falling into the response to stress category, which included classical drought stress markers such as *RD22* and *RAB18* that were already upregulated in basal conditions (Fig 5b). An enrichment of genes belonging to the response to hormone category indicated altered hormonal responses in *BRL3ox* plants under drought (Fig 5a,c and Supplementary Data 7 and Supplementary Data 8). Further analyses of specific hormonal responses revealed that the ABA and jasmonic acid (JA) were the most altered responses (Supplementary Fig. 8). Repression of JA biosynthesis genes may be responsible for decreased levels of JA in basal conditions (Supplementary Fig. 8).

**Figure 5.**
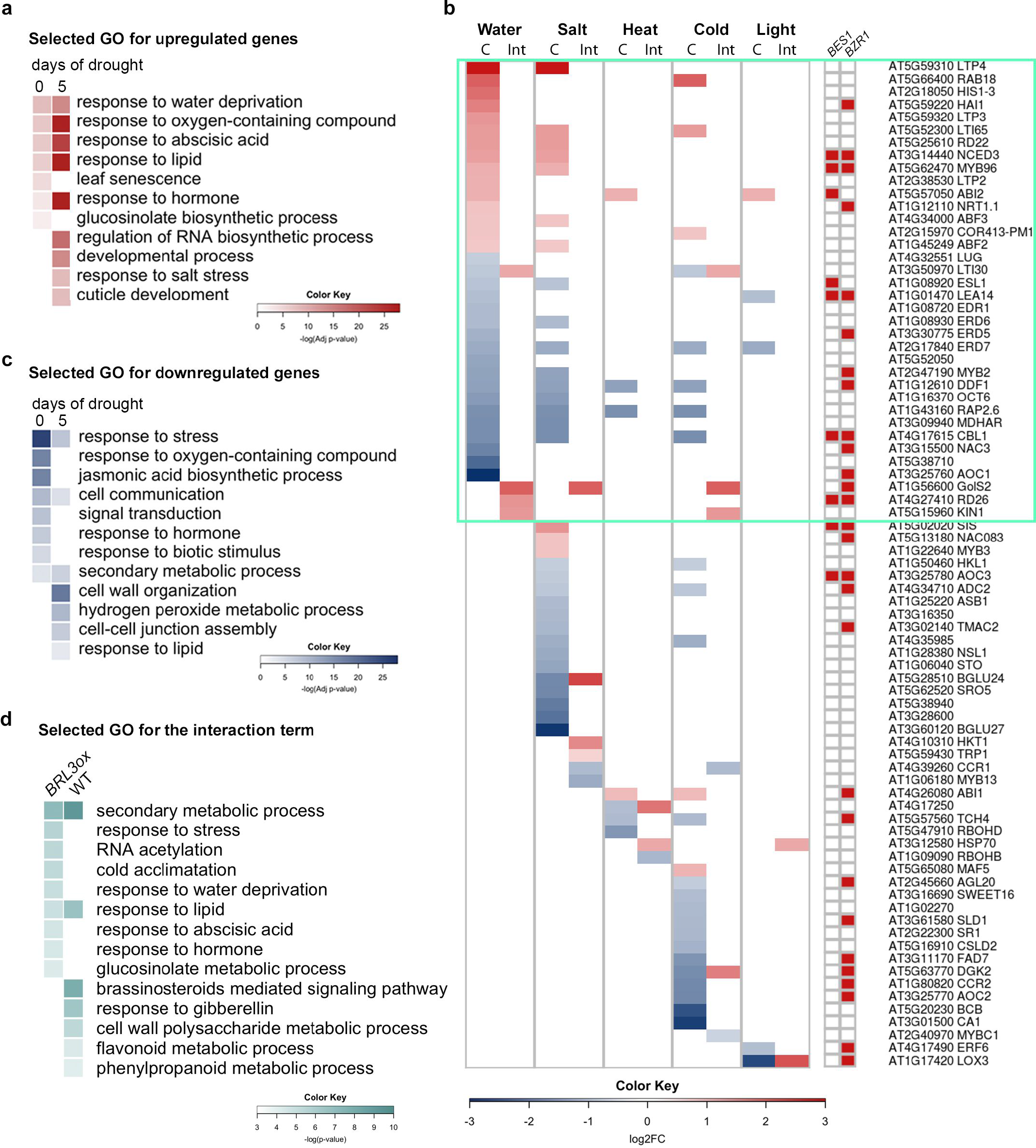
Stress genes are constitutively activated in *BRL3ox* roots. (**a**) Most representative GO categories enriched in *BRL3ox* roots from the upregulated genes at time 0 and after 5 days of drought (**b**) Deployment of genes within “Response to stress” (GO:0006950) term that are also annotated as responsive to water, salt, heat, cold, and light stress. Colors in the heatmap represent the log2 fold change of *BRL3ox* vs. WT roots in control conditions (C) or the differential drought response (log2(FC drought/CTRL in *BRL3ox*)) – (log2(FCdrought/CTRL in WT)) if the gene is affected by the interaction genotype*drought (Int.). Red color in the squared heatmaps on the right shows that the gene has been previously identified as a direct target of BES1 or BZR1 transcription factors. (**c**) Most representative GO categories enriched in *BRL3ox* roots from the upregulated genes at time 0 and after 5 days of drought. (**d**) Most representative GO categories enriched among genes affected by the interaction genotype-drought. GO categories enriched in genes activated in *BRL3ox* under drought compared to WT (left column) in genes repressed in *BRL3ox* under drought compared to WT (right column). Color bars: –log of p-value (adjusted by Benjamini-Hochberg or non-adjusted).

In order to uncover differential drought responses between WT and *BRL3ox* roots, we constructed a linear model accounting for the interaction between genotype and drought (Supplementary Data 9). Taking the 200 most significantly affected genes, we grouped them in (i) genes more activated in *BRL3ox* under drought compared to WT (Supplementary Data 10) and (ii) genes more repressed in *BRL3ox* under drought compared to WT (Supplementary Data 11). GO enrichment analysis of this genotype-drought interaction revealed (i) secondary metabolism, response to stress and response to water deprivation in the first group and (ii) brassinosteroid mediated signaling pathway in the second group (Fig. 5d). Importantly, the expression levels of dehydration response genes remained repressed in *quad* mutant plants during drought (Supplementary Fig. 7 and Supplementary Data 12-15). The expression levels of two key BR biosynthesis genes, *CPD* and *DWF4* were analyzed by RT qPCR. Consistently, within the drought time course, transcription levels of *CPD* and *DWF4* were increased in *quad* and reduced in *BRL3ox* compared to WT plants. Quantification of the bioactive BR hormone Castasterone (CS) showed similar trends and we could only detect BL in *quad*, suggesting that BL is accumulated in *quad* more than in WT and *BRL3ox* plants (Supplementary Fig. 9).

Analysis of transcription factors revealed 29 of them with differential responses to drought between *BRL3ox* and WT roots. Interestingly, the drought-responsive transcription factor *RD26* showed an enhanced response in *BRL3ox* roots during stress, whereas several vascular-specific transcription factors remained repressed under drought (Supplementary Fig. 10). Given that the BRL3 receptor is natively expressed at the phloem-pole pericycle and enriched in vascular tissues when overexpressed ^33^, we analyzed the spatial distribution of the deregulated genes within the root tissues in our RNAseq dataset ^49^. The deregulated genes were enriched for genes that are preferentially expressed in specific vascular tissues such as the pericycle and phloem pole pericycle but also in lateral root primordia (which initiates from pericycle) and root hair cells (Fig. 6a, see Methods). Interaction-affected genes were enriched in pericycle and phloem but also in columella and cortex expressed genes (Fig. 6b). Among the phloem-enriched genes, we found two trehalose phosphatases (*TPPs*) and one galactinol synthases (*GolS2*) that show increased expression in *BRL3ox* roots at basal conditions and in response to drought (Fig. 6d). These enzymes are involved in the synthesis of the osmoprotectant metabolites — trehalose, myo-inositol and raffinose — that overaccumulated in *BRL3ox* roots. Together, these results suggest the importance of changes in expression of phloem-associated genes for sustaining drought resistance.

**Figure 6.**
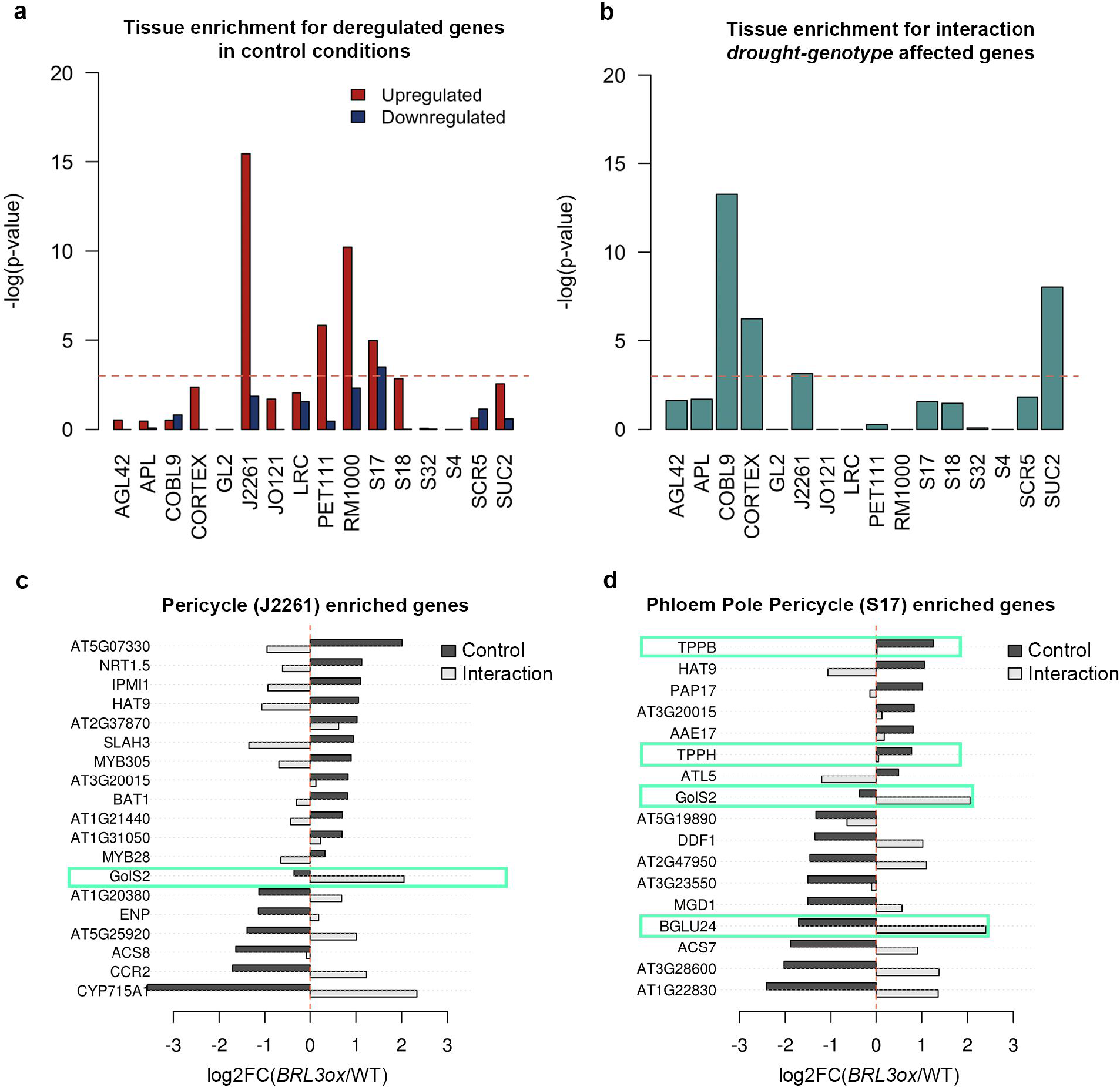
Enrichment of deregulated genes in *BRL3ox* root vasculature. (**a**) Tissue enrichment for upregulated (red) or downregulated (blue) genes in control conditions. Bars trespassing the p-value threshold (0.05) were considered enriched in the dataset. (**b**) Tissue enrichment for genes affected by the interaction genotype*drought Bars trespassing the threshold p-value<0.05 were considered enriched in the dataset. (**a-b**) Deregulated genes tissue enrichment. AGL42: Quiescent center, APL: Phloem + Companion cells, COBL9: Root hair cells, CORTEX: Cortex, GL2: Non-hair cells, J2661: Pericycle, JO121: Xylem pole pericycle, LRC: Lateral root cap, PET111: Columella, RM1000: Lateral root primordia, S17: Phloem pole pericycle, S18: Maturing Xylem, S32: Protophloem, S4: Developing xylem, SCR5: Endodermis, SUC2: Phloem. y-axes represent the negative logarithm of one-tailed Fisher’s test. (**c**) Deregulated genes enriched in the Pericycle (J2261 marker). (**d**) Deregulated genes enriched in the Phloem Pole Pericycle (S17 marker). (**c,d**) Bars represent the log2 fold-change of *BRL3ox* vs. WT roots in control (black) or the difference of drought responses between *BRL3ox* and WT (FC drought/CTRL in *BRL3ox* – FC drought/CTRL in WT) in the lineal model (gray). Blue boxes highlight enzymes directly involved in the metabolism of deregulated metabolites.

Furthermore, a statistical analysis revealed a significant link between the whole transcriptomic and metabolomic signatures, both in basal conditions and under drought (p=0.017 and p=0.001 respectively; see Methods), suggesting that the metabolic signature of *BRL3ox* plants is transcriptionally controlled. We used the metabolic and transcriptomic signatures to identify deregulated metabolic pathways using Paintomics ^50^. This analysis suggests constitutive deregulation of sucrose metabolism in *BRL3ox* plants that was enhanced during drought stress. We also found that BRL3 overexpression affects galactose metabolism under periods of drought, including the raffinose family of oligosaccharides (RFOs) synthesis pathway (Supplementary Fig. 11, Supplementary Data 16 and 17). Collectively, these results suggest that BRL3 overexpression promotes drought tolerance, mainly by controlling sugar metabolism.

## DISCUSSION

Our study shows that overexpression of the BRL3 receptor can prevent growth arrest during drought. We suggest that this is accomplished through the transcriptional control of metabolic pathways that produce osmoprotectant metabolites that accumulate in the roots. While spatial BR signaling has been shown to contribute to stem cell replenishment in response to genotoxic stress ^31,34^, here we show that ectopic expression of vascular-enriched BRL3 receptors can promote growth during drought. Altogether, our results suggest that spatial regulation of BR signaling can affect plant stress responses.

The exogenous application of BR compounds has been used widely in agriculture to extend growth under different abiotic stresses ^22,51^, yet how these molecules precisely activate growth in challenging conditions remains largely unknown. The analysis of BR signaling and BR synthesis mutant plants subjected to stress failed to provide a linear picture of the involvement of BR in drought stress adaptation. For instance, although overexpression of the canonical BRI1 pathway and the BR biosynthesis gene DWF4 can both confer abiotic stress resistance ^24,38^, *BRI1* loss-of-function mutants also showed drought stress resistance ^25,26^. However, increased levels of BR-regulated transcription factors trigger antagonistic effects in drought stress responses ^26,52^, thus depicting a complex scenario for the role of BRs in abiotic stress. Given the spatiotemporal regulation of the BR signaling components ^39^ and the complexity of drought traits ^7^, it is plausible to hypothesize that drought traits are under the control of cell type-specific BR signaling.

Our study unveils that the BR family of receptors, in addition to promoting growth, guides phenotypic adaptation to drought by influencing a myriad of drought stress related traits. The drought resistance phenotypes of BR loss-of-function mutants (Fig. 3a) are likely caused by a reduced exposure of these plants to the effect of drought. This phenomenon, known as drought avoidance, is linked to growth arrest and stress insensitivity that maintains transpiration, leaf water status and photosynthesis along the drought (Fig. 3 and Supplementary Fig. 4). The reduced levels of ABA and canonical stress-related metabolites, together with the downregulation of stress-related genes, further support the insensitivity of *quad* plants to stress (Supplementary Figs. 7 and 8). In contrast, the phenotypes observed in *BRL3ox* plants indicate an active drought-tolerance mechanism driven by overexpression of the BRL3 receptor. First, *BRL3ox* roots showed increased water stress-induced PCD in the root tip compared to WT (Fig. 1c,d), which has been proposed to modify the root system architecture and thereby enhance drought tolerance ^49^. Second, the enhanced hydrotropic response of *BRL3ox* roots (Fig. 2a-c) could function during water-limited conditions by modifying root architecture for increased acquisition of water, favoring plant growth and survival under drought conditions as previously described ^53^. Third, at same RWC in leaves, the rate of photosynthesis and transpiration were more stable in *BRL3ox* than in WT plants during drought (Fig. 3d,e and Supplementary Fig. 4). Altogether, these findings indicate that BRL3 overexpression actively promotes drought tolerance without penalizing plant growth.

We found the expression of the drought-response transcription factor *RD26* to be enhanced in *BRL3ox* roots when subjected to drought (Supplementary Fig. 10). *RD26* has been shown to antagonize the BR canonical transcription factor BES1 ^26^, thereby suggesting that BRL3 overexpression activates alternative pathways. These alternative pathways may be derived from a spatial specialization of BR functions within the root. Indeed, we found that genes preferentially expressed in vascular tissues, especially within phloem-related cell types, were overrepresented among deregulated genes in *BRL3ox* roots (Fig. 6a,b). The localization of the native BRL3 protein in phloem cells ^33^ and the metabolic signature found in *BRL3ox* susuggests a possible role in phloem loading during drought. Moreover, metabolic enzymes implicated in trehalose and RFO metabolism were enriched in vascular tissues and either upregulated in *BRL3ox* roots in basal conditions or strongly responding to drought (Fig. 6c,d). Thus, BRL3 overexpression may affect not only loading and unloading of the phloem, but may also directly control metabolic pathways. This is the case for the trehalose phosphate phosphatase family (*TPPs*) ^54,55^ and galactinol synthase 2 (*GolS2*) ^56^, which are both described to impact drought responses and are involved in trehalose and RFO synthesis respectively. In addition to controlling expression in vascular tissues, our analyses also suggest that BRL3 overexpression regulates non-vascular enzymes important for metabolism and drought responses. These enzymes include hexokinases such as *HXK3* or *HKL1*, the sucrose synthases *SUS3* and *SPS2F*, and proline dehydrogenase genes such as the early response to dehydration 5 (*ERD5)* which is involved in stress tolerance ^57^. In light of our findings and given that *Bes1-D* gain-of-function mutants exhibit drought hypersensitivity ^26^, we propose that overexpression of the vascular BRL3 receptors may act independently of the canonical growth-promoting BRI1 pathway.

Our data further suggest that *BRL3ox* plants accumulate sugars in the sink tissues to enable plant roots to grow and escape drought by searching for water within the soil. In support of these findings, we also observed reduced levels of photosynthesis in well-watered leaves of *BRL3ox* plants (Fig. 3e). These results, together with the higher levels of sucrose in roots compared with in shoots (Fig. 4a) and higher levels of glucose and fructose in the shoots suggest that the BRL3 pathway promotes sugar mobilization from the leaves (source) to the roots (sink). In fact, previous work reported that BRs promote the flow of assimilates in crops from source to sink via the vasculature ^58^ and via sucrose phloem unloading ^59^.

In control conditions, *BRL3ox* plants exhibited a metabolic signature enriched in proline and sugars. Proline and sugar accumulation classically correlates with drought stress tolerance, osmolytes, ROS scavengers, and chaperone functions ^5,45-47,60,61^, suggesting that overexpression of the BRL3 receptor promotes priming ^48,62^. In addition, *BRL3ox* plants also accumulated succinate, fumarate and malate. Importantly, all these metabolites were decreased in *quad* mutant plants. Altogether, these data suggest a role for BRL3 signaling in the promotion of the tricarboxylic acid (TCA) cycle, sugar and amino acid metabolism.

In drought stress conditions, *BRL3ox* shoots displayed increased levels of the amino acids proline, GABA and tyrosine. In contrast, trehalose, sucrose, *myo-*inositol, raffinose, and proline were the most abundant metabolites in the *BRL3ox* roots along the stress time course. Importantly, all these metabolites have previously been linked to drought resistance ^45,46,60^. In addition, the levels of the RFO metabolites raffinose and *myo*-inositol, which are involved in membrane protection and radical scavenging ^63^, were higher in the roots of *BRL3ox* plants under drought, yet reduced in the roots of *quad* plants.

Our data suggest that the roots of *BRL3ox* plants are loaded with osmoprotectant metabolites and are thus better prepared to alleviate drought stress via a phenomenon previously referred to as priming ^48,62^. Altogether these data suggest that drought stress responses are correlated with BRL3 receptor levels in the root vasculature, especially within the phloem, and that this is important for the greater survival rates of *BRL3ox* plants. Future cell type-specific engineering of signaling cascades stands out as a promising strategy to circumvent growth arrest caused by drought stress.

## METHODS

### Plant materials

Seeds were sterilized with 35% NaClO for 5 min and washed five times for 5min with sterile dH_2_O. Sterile seeds were vernalized 48 h at 4°C and grown in half-strength agar Murashige and Skoog (MS1/2) media with vitamins and without sucrose. Plates were grown vertically in long day (LD) conditions (16 h of light / 8 h of dark; 22°C, 60% relative humidity). Genotypes used in this study: Columbia-0 WT (Col-0 WT), *brl1-1brl3-1* (*brl1brl3*), *bak1-3* (*bak1*), *bri1-301* (*bri1*), *bri1-301brl1-1brl3-1* (*bri1brl1brl3*), *bri1-301bak1-3brl1-1brl3-1 (quad)* and *35S:BRL3-GFP (BRL3ox)* ^33^. DNA rapid extraction protocol ^64^ was used for all the plant genotyping experiments. Supplementary Table 2 describes the primers used for genotyping of the BR mutant plants.

### Brassinolide and sorbitol sensitivity assays in roots

For hormone treatments, seeds were continuously grown in concentration series of brassinolide (BL,Wako, Japan). For sorbitol assays, three-day-old seedlings were transferred to either control or 270 mM sorbitol media for four additional days. The root length of seven-day-old seedlings was measured using Image J (http://rsb.info.nih.gov/ij/) and compared with automatically acquired data from the MyROOT ^65^ software (Supplementary Fig. 12). Four-dayold roots grown in control conditions or in 24 h of sorbitol were stained with propidium iodide (10 ug/ml, PI, Sigma). PI stains the cell wall (control) and DNA in the nuclei upon cell death (sorbitol). Images were acquired with a confocal microscope (FV1000 Olympus). Cell death damage in primary roots was measured in a window of 500 µm from QC in the middle root longitudinal section (Image J). As an arbitrary setting to measure the stained area, a color threshold ranging from 160 to 255 in brightness was selected.

### Root hydrotropism

Seedlings were germinated in MS1/2 without sucrose for six days. Then, the lower part of the agar was removed from the plates and MS1/2 with 270 mM sorbitol was added to simulate a situation of reduced water availability. The media was placed in 45-degree angle to scape gravitropism effect. When indicated, 1 μM of brassinazole ^44^ was added to sorbitol media. Root curvature angles were measured and analyzed using the Image J software (http://rsb.info.nih.gov/ij/).

### Drought stress for scoring plant survival

One-week-old seedlings grown in MS1/2 agar plates were transferred individually to pots containing 30±1 g of substrate (plus 1:8 v/v vermiculite and 1:8 v/v perlite). For each biological replicate, 40 plants of each genotype were grown in LD conditions for three weeks. Three-week-old plants were subjected to severe drought stress by withholding water for 12 days followed by re-watering. After the seven-day recovery period, the surviving plants were photographed and manually counted (two-sided chi-squared test, p-value <0.01).

### Metabolite profiling analyses

One-week-old seedlings were placed in individual pots with 30 g of autoclaved soil and grown under LD photoperiodic conditions. After three weeks growing, half of the plants were subjected to severe drought (withholding water) for six days and the other half were watered normally (well-watered control conditions). A total of five biological replicates were collected every 24 h during the time course (from day 0 to day 6) both in drought and watered conditions and for each genotype (WT, *quad* and *BRL3ox*). Four independent plants were bulked in each biological replicate. Roots were manually separated from shoots. Four entire shoots were grinded using the Frosty Cryogenic grinder system (Labman). Four entire root samples were grinded in the Tissue Lyser Mixer-Mill (Qiagen). Roots were aliquoted into 20 mg samples and shoot into 50 mg samples (the exact weight was annotated for data normalization). Primary metabolite extraction was carried as follows ^66^. One Zirconia and 500 μl of 100% Methanol premixed with Ribitol (20:1) were added and samples were subsequently homogenized in the Tissue Lyser (Qiagen) 3 min at 25 Hz. Samples were centrifuged 10 min at 14000 rpm (10 °C) and resulting supernatant was transferred into fresh tubes. Addition of 200 μl of CHCl3 and vortex ensuring one single phase followed by the addition of 600 μl of H20 and vortex 15 sec. Samples were centrifuged 10 min at 14000 rpm (10 °C). 100 μl from the upper phase (polar phase) were transferred into fresh eppendorf tubes (1.5 ml) and dried in the speed vacuum for at least 3 h without heating. 40 μl of derivatization agent (methoxyaminhydrochloride in pyridine) were added to each sample (20 mg/ml). Samples were shaken during 3 h at 900 rpm at 37 °C. Drops on the cover were shortly spun down. One sample vial with 1 mL MSTFA + 20 μl FAME mix was prepared. Addition of 70 μl MSTFA + FAMEs in each sample was done followed by shaking 30 min at 37 °C. Drops on the cover were shortly spun down.

Samples were transferred into glass vials specific for injection in GC-TOF-MS. The GC-TOF-MS system comprised of a CTC CombiPAL autosampler, an Agilent 6890N gas chromatograph, and a LECO Pegasus III TOF-MS running in EI+ mode. Metabolites were identified by comparing to database entries of authentic standards ^67^. Chromatograms were evaluated using Chroma TOF 1.0 (Leco) Pegasus software was used for peak identification and correction of RT. Mass spectra were evaluated using the TagFinder 4.0 software ^68^ for metabolite annotation and quantification (peak area measurements). The resulting data matrix was normalized using an internal standard, Ribitol, in 100% methanol (20:1), followed by normalization with the fresh weight of each sample. Metabolomics data from control (well-watered) conditions at day 0 were analyzed with a two-tailed t-test, p-value<0.05 (no multiple testing correction). Data from the time course was analyzed with R software using the maSigPro package ^69^. Briefly, the profile of each metabolite under each condition was fitted to a polynomial model of maximum degree 3. The curves of each genotype were statistically compared taking into account the fitting value and correcting the p-value (Benjamini-Hochberg method). Significant metabolites (p-value < 0.05) having a differential profile between genotypes were plotted to visualize their behavior under the drought time course. Clustering analysis was performed using the maSigPro package and the *hclust* R core function.

### Transcriptomic profiling analysis

For microarray analysis, a drought stress time course was carried out in WT and *quad* mutant three-week-old plants. Entire plants grown under drought stress and control conditions were collected every 48 h during the time course (Day 0, Day 2 and Day 4). Two biological replicates composed of five independent rosettes were collected. RNA was extracted with the Plant Easy Mini Kit (Qiagen) and quality checked using the Bioanalyser. A Genome-Wide Microarray platform (Dual color, Agilent) was performed by swapping the color hybridization of each biological replicate (Cy3 and Cy5). Statistical analysis was performed with the package “limma” ^70^, and the “mle2/“normexp” background correction method was used. Different microarrays were quantile-normalized and a Bayes test used to identify differentially expressed probes. The results were filtered for adjusted p-value<0.05 (after Benjamini-Hochberg correction) and Log2 FC >|1.5|. For RNAseq analysis, three-week-old roots were detached from mature plants grown in soil under control conditions and five days of drought. RNA was extracted as described above. Stranded cDNA libraries were prepared with TruSeq Stranded mRNA kit (Illumina). Single-end sequencing, with 50-bp reads, was performed in an Illumina HiSeq500 sequencer, at a minimum sequencing depth of 21 M. Reads were trimmed 5 bp at their 3’ end, quality filtered and then mapped against the TAIR10 genome with “HISAT2”. Mapped reads were quantified at the gene level with “HtSeq”. For differential expression, samples were TMM normalized and statistical values calculated with the “EdgeR” package in R. Results were filtered for adjusted p-value (FDR) <0.05 and FC >|2| in the pairwise comparisons. For the evaluation of differential drought response between WT and *BRL3ox* roots, a lineal model accounting for the interaction genotype and drought was constructed with “EdgeR” package. The interaction term was evaluated. A gene was considered to be affected by the interaction if p-value (uncorrected) < 0.0025. Heatmaps were performed in R with the heatmap.2 function implemented in the “gplots” package.

For the Rt qPCR, cDNA was obtained from RNA samples by using the Transcriptor First Strand cDNA Synthesis Kit (Roche) with oligo dT primers. qPCR amplifications were performed from 10ng of cDNA using LightCycler 480 SYBR Green I master mix (Roche) in 96-well plates according the manufacturer recommendations. The Real Time PCR was performed on a LightCycler 480 System (Roche). Ubiquitin (AT5G56150) was used as housekeeping gene for relativizing expression. Primers used are described in Supplementary Table 3.

### Statistical methods and omics data integration

For root tissue enrichment analysis, deregulated genes were queried against available lists of tissue-enriched genes ^49^. For each tissue, a 2×2 contingency table was constructed, counting the number of deregulated genes in the tissue that were enriched and non-enriched and also the number of non-deregulated genes (for either FDR>0.05 or logFC >/< in the RNAseq gene universe) that were enriched and non-enriched. Statistical values of the enrichment were obtained using a one-sided Fisher’s test. To statistically evaluate the influence of transcriptomic changes on the metabolic signature, both deregulated enzymes and metabolites were queried in an annotation file of the metabolic reactions of *Arabidopsis thaliana*, which included merged data from the KEGG (http://www.genome.jp/kegg/) and BRENDA (www.brenda-enzymes.org) databases. Then, the same approach of constructing a 2×2 contingency table was taken. Significant and non-significant metabolites annotated in the database were matched with differentially and non-differentially expressed genes annotated in the database. The statistical value of the association between regulated metabolites and genes was obtained through a two-sided Fisher’s exact test. Genes and metabolites were mapped onto the KEGG pathways using the PaintOmics3 (http://bioinfo.cipf.es/paintomics/) according the developer’s instructions ^50^.

### Physiological parameters and chlorophyll fluorescence

One-week-old seedlings were placed in individual pots and watered with the same volume of a modified Hoagland solution (one-fifth strength). Pots were weighed daily during the experiment. Well-watered control plants were grown in 100% field capacity (0% of water loss). The time course drought stress assay was started by withholding the nutrient solution until reaching 25, 50, 60 and 70% water loss. Photosynthesis (*A*) and transpiration (*E*) were measured in control and drought plants at those time points. Four plants of each genotype were harvested at 0, 50 and 70% water loss for biomass, water content and hormone analyses. Drought experiments were repeated three times and at least four plants per genotype and treatment were used in each experiment. RWC was calculated according to the formula: RWC (%) = [(FW-DW) / (TW-DW)] x 100.

### Plant hormones quantification

Plant hormones cytokinins (*trans*-zeatin), gibberellins (GA1, GA4 and GA3), indole-3-acetic acid (IAA), abscisic acid (ABA), salicylic acid (SA), jasmonic acid (JA), and the ethylene precursor 1-aminocyclopropane-1-carboxylic acid (ACC) were analyzed as follows. 10 μl of extracted sample were injected in a UHPLC– MS system consisting of an Accela Series U-HPLC (ThermoFisher Scientific, Waltham, MA, USA) coupled to an Exactive mass spectrometer (ThermoFisher Scientific, Waltham, MA, USA) using a heated electrospray ionization (HESI) interface. Mass spectra were obtained using the Xcalibur software version 2.2 (ThermoFisher Scientific, Waltham, MA, USA). For quantification, calibration curves were constructed for each analyzed hormone (1, 10, 50, and 100 μg l−1) and corrected for 10 μg l−1 deuterated internal standards. Recovery percentages ranged between 92 and 95%.

For endogenous BR analysis plant materials (4 g fresh weight) were lyophilized and grinded. BL and CS were extracted with methanol and purified by solvent partitions by using a silica gel column and ODS-HPLC as follows. The endogenous levels of BL and CS were quantified by LC-MS/MS using their deuterated internal standards (2ng).

LC-MS/MS analysis was performed with a triple quadrupole/linear ion trap instrument (QTRAP5500; AB Sciex, USA) with an electrospray source. Ion source was maintained at 300_ᄎ_C. Ion spray voltage was set at 4500 V in positive ion mode. MRM analysis were performed at the transitions of m/z 487 to 433 (Collision Energy, CE 30 V) and 487 to 451 (CE 21 V) for 2 H 6 -BL, m/z 481 to 427 (CE 30 V) and 481 to 445 (CE 30 V) for BL, m/z 471 to 435 (CE 23 V) and 471 to 453 (CE 25 V) for 2 H 6 -CS and m/z 465 to 429 (CE 23 V) and 465 to 447 (CE 25 V) for CS. Enhanced product ion scan was carried out at CE 21 V. HPLC separation was performed using a UHPLC (Nexera X2; Shimadzu, Japan) equipped with an ODS column (Kinetex C18, f2.1 ‘ 150 mm, 1.7 μm; Phenomenex, USA). The column oven temperature was maintained at 30°C. The mobile phase consisted of acetonitrile (solvent A) and water (solvent B), both of which contained 0.1% (v/v) acetic acid. HPLC separation was conducted with the following gradient at flow rate of 0.2 mL·min-1: 0 to 12 min, 20% A to 80% A; 12 to 13 min, 80% A to 100% A; 13 to 16 min, 100% A.

## DATA AVAILABILITY

RNAseq and microarray data that support the findings of this study have been deposited in Gene Expression Omnibus (GEO) with the GSE119382 [https://www.ncbi.nlm.nih.gov/geo/query/acc.cgi?acc=GSE119382”] and GSE119383 [https://www.ncbi.nlm.nih.gov/geo/query/acc.cgi?acc=GSE119383] accession codes.

## ACKNOWLEDGEMENTS

We thank Riccardo Aiese Cigliano (Sequentia Biotech) for help with microarray analysis, Tony Ferrar for critical manuscript revision and language editing (http://www.theeditorsite.com) and Olga Moreno Pradas for graphic design support. We acknowledge financial support from the Spanish Ministry of Economy and Competitiveness, through the “Severo Ochoa Programme for Centres of Excellence in R&D” 2016-2019 (SEV-2015-0533)”, the CERCA Programme from the Generalitat de Catalunya, and the Agencia de Gestió d’Ajuts Universitaris i de Recerca (2014 SGR 1406). N.F. was funded by Fundación Renta Corporación, by EMBO short-term postdoctoral fellowship (ASTF 422-2015) and by BIO2013-43873; F.L.E was funded by BIO2013-43873 and BIO2016-78150-P; and D.B. was funded by BIO2013-43873 and ERC-2015-CoG-GA 683163. CMA, AA and FPA acknowledge financial support from the Spanish MINECO-FEDER (project AGL2014-59728-R) and from the European Union’s Seventh Framework Programme for research technological development and demonstration under grant agreement no. 289365 (project ROOTOPOWER). S.O. was supported by Ministry of Science and Innovation and University of Malaga (Spain) through the grant Ramón y Cajal program (Sonia Osorio, RYC-09170). M.B was funded by BES-2012-053274 and J.L.R laboratory is supported by grant BFU2014-58289-P from the Spanish Ministry of Economy and Competitiveness. A.I.C.-D. – A.C. collaboration was funded by the European Regional Development Funds and Marie Curie IRSES Project DEANN (PIRSES-GA-2013-612583). A.I.C.-D. is a recipient of BIO2013-43873 and BIO2016-78150-P grants from the Spanish Ministry of Economy and Competitiveness. A.I.C.-D. is recipient of an ERC Consolidator Grant; this project has received funding from the European Research Council (ERC) under the European Union’s Horizon 2020 research and innovation program (grant agreement No 683163).

## AUTHOR CONTRIBUTIONS

A.I.C.-D. conceived, designed and supervised the study. N.F., F.L.E, D.B., C.M.A. and F.P.A. performed the genetics and stress phenomic experiments and data analyses. N.F., F.L.E., M.B., J.L.R. and A.I.C.-D. designed the genome-wide experiments. N.F., F.L.E., A.C. and A.I.C.-D. performed the genome-wide experiments and data analysis. N.F., F.L.E., T.T., S.O., and A.R.F. carried out the metabolic profiling and data analysis. A.A. performed the hormone profiling experiments. T.N. and T.Y. carried the CS and BL quantification assays. N.F., F.L.E and A.I.C.-D. wrote the manuscript. All authors discussed the results and the manuscript.

